## Supplementary material for "Quantitative analysis questions the role of MeCP2 as a global regulator of alternative splicing": S1 Appendix

### S1 Appendix. Detailed information about software, software versions and command line arguments used for analysis

#### DNA methylation regions & ChIP-seq signal regions

Region specific mean methylation, methylation density and ChIP signal are calculated for the regions described in S1 Appendix Fig 1

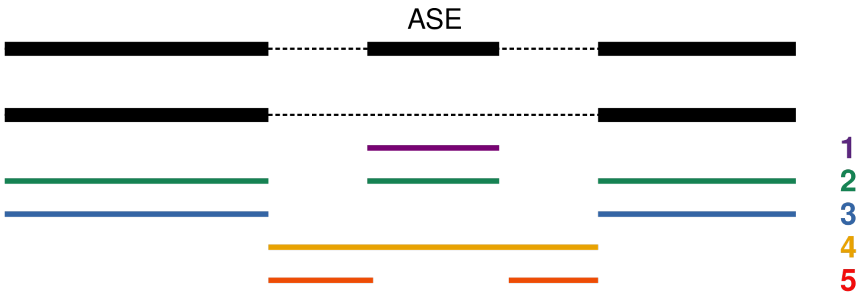


**S1 Appendix Fig 1.** Regions of gene for which DNA methylation metrics are calculated 1) ASE (alternative spliced exon) 2) combination of all exons containing the ASE 3) combination of all exons without the ASE 4) combination of all introns 5) combination of all introns without the ASE.

#### Genome assembly and annotation

*Homo sapiens* assembly (hg19) is used for the GSE125660 [?] and *Mus musculus* assmebly (mm10) is used for GSE128186 [?], GSE103214 [?], GSE48307 [?] and GSE85517 [?]. The genome annotation for alternative splicing events are made available as part of BRIE [?] and downloaded from `https://sourceforge.net/projects/brie-rna/files/annotation/`. HISAT2 [?] genome indexes for hg19 and mm10 are downloaded from `ftp://ftp.ccb.jhu.edu/pub/infphilo/hisat2/data`.

#### Software and command-line arguments

After downloading the raw reads from Gene Expression Omnibus archive, we used HISAT2 [?] v2.0.2 for aligning the reads to the genome using the following command.

```
hisat2 -x {genome.index} -1 {input.one} -2 {input.two} --dta-cufflinks --known-splicesite-
    infile {splicesite.file} -k 2 --no-unal -p 4 | samtools view -h -b -@ 4 -o {output} -
```

Miso [?] v0.5.4 is used to quantify splicing ratios from the aligned files. The `indexed.assembly` is generated from genome annotation for alternative splicing events using the command `index_gff` which is part of Miso.

```
miso --run {indexed.assembly} {file.bam} --output-dir {outdir} --paired-end {mean.fragment.
    length} {stdev.fragment.length} --settings-filename miso_settings.txt --read-len {read.
    length}
```

Miso [?] v0.5.4 is used to estimate differential splicing between samples. Alignments from different replicates are pooled into a single BAM file and then used for comparison.

```
compare_miso --compare-samples {treatment} {control} {outdir}
```

BRIE [?] v0.2.0 is used to quantify splicing ratios from the aligned files.

```
brie -a {gtf} -s {file.bam} -f {brie.features} -o {outdir}
```

Cufflinks [?] v2.2.1 is used to quantify splicing ratios from the aligned files.

```
cufflinks -G {gtf} -o {outdir} {file.bam}
```

Differential gene expression is estimated using DESeq2 [?] v1.24.0 with genotype or age as the covariate in the `design` formula.

#### Generating Simulation Data

Sequencing reads were generated using Art https://www.niehs.nih.gov/research/resources/software/biostatistics/art/index.cfm on reference RNA sequences from Sequins https://www.sequinstandards.com at different coverage depths.

```
art_illumina -na -i RNAsequins.v2.2.fa -l 100 -ss HS25 -f 25 -m 200 -s 10 -o fastq/sim1.
art_illumina -na -i RNAsequins.v2.2.fa -l 100 -ss HS25 -f 50 -m 200 -s 10 -o fastq/sim2.
art_illumina -na -i RNAsequins.v2.2.fa -l 100 -ss HS25 -f 100 -m 200 -s 10 -o fastq/sim3.
```

Reads were then aligned using HISAT2 [?] as described earlier and Splicing Ratios were quantified using Miso [?] and Cufflinks [?]. Splicing Ratios were then visualized as in S3 Fig D.

#### Regressing Sequence Features with Splicing Ratios

The modelling with Sequence Features and differences in DNA methylation patterns with differences in Splicing Ratios is executed using the same way as described below.

```
from scipy import stats
from sklearn.model_selection import cross_validate
from sklearn import linear_model
from sklearn import ensemble

import glob
import pandas as pd

miso_summaries = glob.glob("*.miso_summary")
features = pd.read_csv("factors.csv.gz", compression="gzip", index_col=0)

r2_scores = []

for summary in miso_summaries:
        sample = summary.split(".miso_summary")[0]
        psi = pd.read_csv(fraction, index_col=0, usecols=[0,1], sep="\t").replace(0,0.00001).
            replace(1, 0.99999).apply(stats.norm.ppf)
        y = psi["miso_posterior_mean"]
        X = features.loc[y.index]

        lm = linear_model.BayesianRidge(n_iter=1000, tol=0.00001, normalize=True,
            compute_score=True)
```

```
        lm_scores = cross_validate(lm, X, y, cv=5, scoring=('r2'), n_jobs=-1)
        for score in lm_scores["test_scores"]:
                r2_scores.append(["Linear␣Regression", "Miso", sample, score])

        rf = ensemble.RandomForestRegressor(n_estimators=100, random_state=0, n_jobs=-1)
        rf_scores = cross_validate(rf, X, y, cv=5, scoring=('r2'), n_jobs=-1)
        for score in rf_scores["test_scores"]:
                r2_scores.append(["Random␣Forest␣Regression", "Miso", sample, score])

brie_fractions = glob.glob("*.fractions.tsv")

for fraction in brie_fractions:
        sample = summary.split(".fractions.tsv")[0]
        psi = pd.read_csv(fraction, index_col=0, sep="\t", usecols=[0,5]).replace(0,0.00001).
            replace(1, 0.99999).apply(stats.norm.ppf)
        X = features.copy()
        X.index = ["{}.in".format(ii) for ii in X.index]
        y = psi.loc[X.index]["Psi"]

        lm = linear_model.BayesianRidge(n_iter=1000, tol=0.00001, normalize=True,
            compute_score=True)
        lm_scores = cross_validate(lm, X, y, cv=5, scoring=('r2'), n_jobs=-1)
        for score in lm_scores["test_scores"]:
                r2_scores.append(["Linear␣Regression", "BRIE", sample, score])

        rf = ensemble.RandomForestRegressor(n_estimators=100, random_state=0, n_jobs=-1)
        rf_scores = cross_validate(rf, X, y, cv=5, scoring=('r2'), n_jobs=-1)
        for score in rf_scores["test_scores"]:
                r2_scores.append(["Random␣Forest␣Regression", "BRIE", sample, score])
```

#### Scatter Plot

```
import numpy as np
import pandas as pd
import matplotlib.pyplot as plt
from scipy.stats import gaussian_kde

df = pd.read_csv("y_test_vs_y_pred.tsv", sep="\t", index_col=0)
fig, ax = plt.subplots()
y_te = df["y_test"]
y_pr = df["y_pred"]
xy = np.vstack([y_te, y_pr])
z = gaussian_kde(xy)(xy)
ax.scatter(y_te, y_pr, s=30, c=z, edgecolor='', cmap="Greys")
ax.set_xlim(-3, 3)
ax.set_ylim(-3, 3)
ax.set_xticks([-2, 0, 2])
ax.set_yticks([-2, 0, 2])
ax.plot(bx.get_xlim(), bx.get_ylim(), ls="--", c=".3")
```

#### Regression Analysis Overview

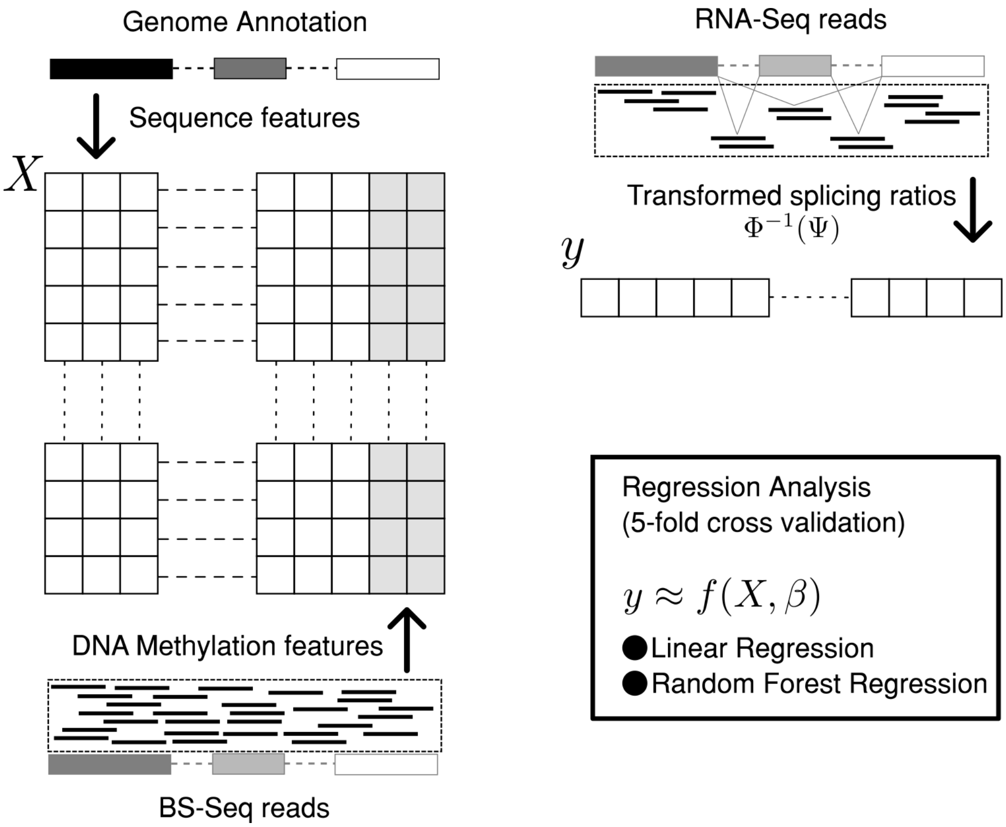


**S1 Appendix Fig 2.** Graphical representation of the overall regression analyses. Accumulation of sequence features from genome annotation, accumulation of DNA Methylation features from BS-Seq (Bisulfite Sequencing) reads and quantification of Splicing Ratios ($\Psi$) from RNA-Seq (RNA Sequencing) reads.
